# The KDEL Trafficking Receptor Exploits pH to Tune the Strength of an Unusual Short Hydrogen Bond

**DOI:** 10.1101/2020.07.18.209858

**Authors:** Zhiyi Wu, Simon Newstead, Philip C. Biggin

**Affiliations:** Department of Biochemistry, South Parks Road, Oxford. OX1 3QU. UK

**Keywords:** transporter, chaperone, molecular simulation, quantum mechanics, protonation

## Abstract

The endoplasmic reticulum (ER) is the main site of protein synthesis in eukaryotic cells and requires a high concentration of luminal chaperones to function. During protein synthesis, ER luminal chaperones are swept along the secretory pathway and must be retrieved to maintain cell viability. ER protein retrieval is achieved by the KDEL receptor, which recognises a C-terminal Lys-Asp-Glu-Leu (KDEL) sequence. Recognition of ER proteins by the KDEL receptor is pH dependent, with binding occurring under acidic conditions in the Golgi and release under conditions of higher pH in the ER. Recent crystal structures of the KDEL receptor in the apo and peptide bound state suggested that peptide binding drives the formation of a short-hydrogen bond that locks the KDEL sequence in the receptor and activates the receptor for COPI binding in the cytoplasm. Using quantum mechanical calculations we demonstrate that the strength of this short hydrogen bond is reinforced following protonation of a nearby histidine, linking receptor protonation to high affinity peptide binding. Protonation also controls the wetting of a cavity adjacent to the peptide binding site, leading to a conformational change that ultimately allows the complex to be recognized by the COPI system.

## Introduction

In eukaryotic cells newly synthesised proteins destined for secretion pass from the endoplasmic reticulum (ER) to the Golgi ^1^. Luminal ER chaperones and quality control enzymes are normally associated with these proteins and are trafficked along with their substrates to the Golgi apparatus. To maintain the required pool of folding chaperones in the ER, it is essential that these proteins are retrieved ^2^. ER protein retrieval is mediated by the KDEL receptor (KDELR), which recognises escaped proteins in the Golgi and mediates their return to the ER via COPI vesicles ^3,4^. The KDELR recognizes escaped ER proteins through a C-terminal ER retrieval sequence (ERS), consisting of Lys-Asp-Glu-Leu (KDEL) ^5,6^, although variations to the canonical sequence exist ^7,8^. Binding of the KDEL sequence to the receptor is pH dependent ^9^. In the acidic environment of the Golgi lumen the receptor forms a stable complex with the KDEL bearing cargo proteins. The activated receptor then signals across the membrane to recruit COPI and initiate retrograde trafficking to the ER. Following retrieval, the higher pH in the ER results in deprotonation of the receptor and release of the KDEL peptide and associated cargo protein, whereupon the receptor is cycled back to the Golgi via COPII vesicles ^10,11^.

Although it was discovered over two decades ago, the molecular mechanism of pH-dependent binding by the KDELR remains stubbornly elusive ^12^. Recently the crystal structure of the KDELR has been solved in both KDEL bound (pH 6.0) and Apo state (pH 9.0) ^13^ (**Fig. 1**). The structures revealed the conformational changes that occur upon signal peptide binding, which alter the electrostatic surface of the receptor on the cytoplasmic side of the membrane that likely mediates the interaction of the receptor with either COPI or COPII to allow for KDEL dependent recycling of the receptor within the secretory pathway. *In vitro* binding and cellular retrieval assays highlighted an essential role for a conserved histidine (H12) at the base of the signal peptide binding pocket. Due to the protonatable nature of this residue within the physiological range of the secretory pathway, it was proposed that this side chain may form the pH sensor for the KDEL retrieval system. This histidine is located adjacent to a short hydrogen bond formed between a conserved tyrosine (Y158) and glutamate (E127), which functions to lock the receptor in an active state following binding of the KDEL ERS. But how exactly does the protonation state of H12 influence the energetics of the system in a way that reflects and explains the functional cycle of the KDEL receptor?

**Figure 1.**
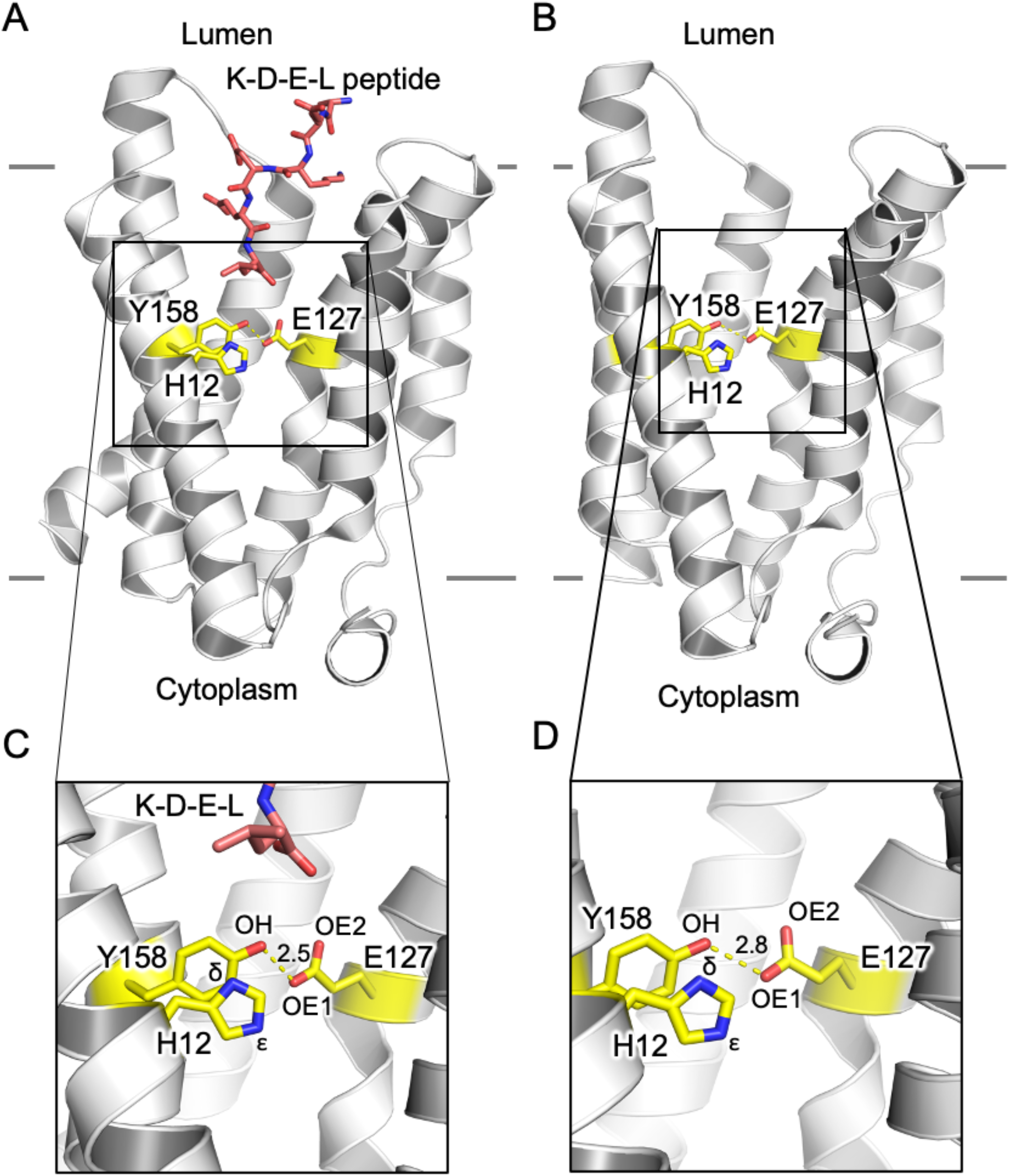
Structures of the KDEL receptor used in this study. (A) and (C) KDEL peptide bound (PDB: 6i6h) and (B, D) the Apo (PDB: 6i6b) state. The hydrogen bond between E127 and Y158 is slightly shorter when peptide is bound at 2.5 Å (C) compared to 2.8 Å in the apo form (D). Atom labels used within this study are also shown along with the position of H12, which has previously be shown to act as the pH sensor. The approximate location of the membrane is shown with solid gray lines.

Using a combination of molecular dynamics (MD) and quantum mechanics (QM) calculations we explored the role of this conserved histidine side chain in pH sensing, and the influence this has on the physical properties of the short hydrogen bond located at the core of the receptor. Our results show that protonation of H12 is essential for the formation of the short hydrogen bond, and thus plays an essential role in stabilising the active state of the receptor. We also show that the presence of this short hydrogen-bond appears to dictate whether a continuous water cavity can form, which our simulations suggest may play an important role in the structural changes that result in signaling the recruitment of either the COPI or COPII coatomer proteins.

## Results

### Establishing the position of the proton

It is well known that X-ray diffraction experiments often cannot resolve the precise location of hydrogen atoms and indeed for this reason they are omitted from the refined models. To characterize the nature of the hydrogen bond between E127 and Y158 and the influence of the nearby H12, we first needed to obtain precise positions of the hydrogen that would, under most conditions, be strongly associated with the hydroxyl oxygen of the tyrosine. Since, in this case our working hypothesis is that there is a particularly short hydrogen bond between the E127 and Y158, we do not know, *a priori*, which atom the hydrogen bond could be bonded to and there are three candidate oxygen atoms to consider; the hydroxyl oxygen of Y158 (labelled OH), and the two carboxy oxygens of E157 (labelled OE1 and OE2, see **Fig. 1**). We thus set up geometry optimizations (**see Methods**) where a hydrogen atom is initially positioned next to one of these oxygens. This approach at least allows the coordinates to adopt a “correct from the quantum mechanical point of view” position of the hydrogens. In the Apo receptor, all hydrogens remain close to their original position after geometry optimization. In the KDEL-bound conformation, when the hydrogen is placed next to E127:OE1, which is the oxygen of the E157 closest to Y158, the position of the hydrogen converges during the optimization toward the oxygen of Y158 and indeed overlaps with the geometry optimized position of the hydrogen when it is initially placed next to the oxygen of Y158 (Y158:OH). Contrary to this, if the hydrogen is positioned closed to E127:OE2, the hydrogen remains bound to that oxygen. Thus, there are clear differences in the energetic landscape of the hydrogen in these different conformations.

To predict the most probable location of the hydrogen among these three oxygens, we computed the free energy of the geometry optimised structures. As is shown in **Table 1**, protonated Y158 in complex with the deprotonated E127 (E127···H—Y158) always has the lowest free energy compared to the two protonated forms of E127 (E127—HE1···Y158 and E127—HE2···Y158). To understand this further, we computed the surface (0.001 a.u. ^14^) electrostatic potentials. As is shown in **Fig. 2**, only E127···H—Y158 shows good charge complementary, whereas both E127—HE1···Y158 and E127—HE2···Y158 have significant overlap between negative charge, which render them energetically less favourable.

**Table 1.** QM characterization of the E127:Y158 hydrogen bond of the KDEL receptor.

|  |  | Free energy<br>(Hartrees) | R(D-H) | R(H...A) | A(D-H...A) |
| --- | --- | --- | --- | --- | --- |
| **KDEL-bound** |  |  |  |  |  |
| (HID12) | E127...H—Y158 | -2245.33 | 1.044 | 1.496 | 174.264 |
| (HID12) | E127—HE2...Y158 | -2245.26 | 0.995 | 2.150 | 145.370 |
| (HIE12) | E127...H—Y158 | -2245.33 | 1.029 | 1.510 | 174.352 |
| (HIE12) | E127—HE2...Y158 | -2245.25 | 0.997 | 2.148 | 145.528 |
| (HIP12) | E127...H—Y158 | -2245.79 | 1.058 | 1.482 | 173.371 |
| (HIP12) | E127—HE2...Y158 | -2245.74 | 0.991 | 2.166 | 144.051 |
| **Apo** |  |  |  |  |  |
| (HID12) | E127...H—Y158 | -2245.33 | 1.004 | 1.861 | 163.691 |
| (HID12) | E127—HE1...Y158 | -2245.31 | 1.039 | 1.803 | 174.172 |
| (HID12) | E127—HE2...Y158 | -2245.27 | 0.985 | 2.206 | 145.279 |
| (HIE12) | E127...H—Y158 | -2245.33 | 0.998 | 1.868 | 163.505 |
| (HIE12) | E127—HE1...Y158 | -2245.31 | 1.052 | 1.790 | 174.668 |
| (HIE12) | E127—HE2...Y158 | -2245.26 | 0.987 | 2.205 | 145.376 |
| (HIP12) | E127...H—Y158 | -2245.81 | 1.005 | 1.867 | 161.615 |
| (HIP12) | E127—HE1...Y158 | -2245.79 | 1.034 | 1.812 | 171.474 |
| (HIP12) | E127—HE2...Y158 | -2245.76 | 0.983 | 2.224 | 143.936 |

Free energy is computed at the level of D3LYP-D4/def2-TZVP. R(D-H) is the intramolecular distance between the donor heavy atom and the hydrogen. R(D-H) is the intramolecular distance between the donor heavy atom and the hydrogen. R(H...A) is the intermolecular distance between the hydrogen and the acceptor heavy atom. A(D-H...A) is the angle of the hydrogen bond along the donor heavy atom, the hydrogen and the acceptor heavy atom. HID12 indicates the system is optimised under a histidine which was protonated on the delta nitrogen, whereas HIE12 indicates that the histidine was protonated on the epsilon nitrogen. HIP12 means that both nitrogens were protonated. E127...H—Y158 stands for the hydrogen bond system between protonated Y158 and deprotonated E127, while E127—HE1...Y158 and E127—HE2...Y158 signifies the deprotonated Y158 hydrogen bonding to the E127 protonated on the E127:OE1 and E127:OE2 oxygen atoms respectively.

**Figure 2.**
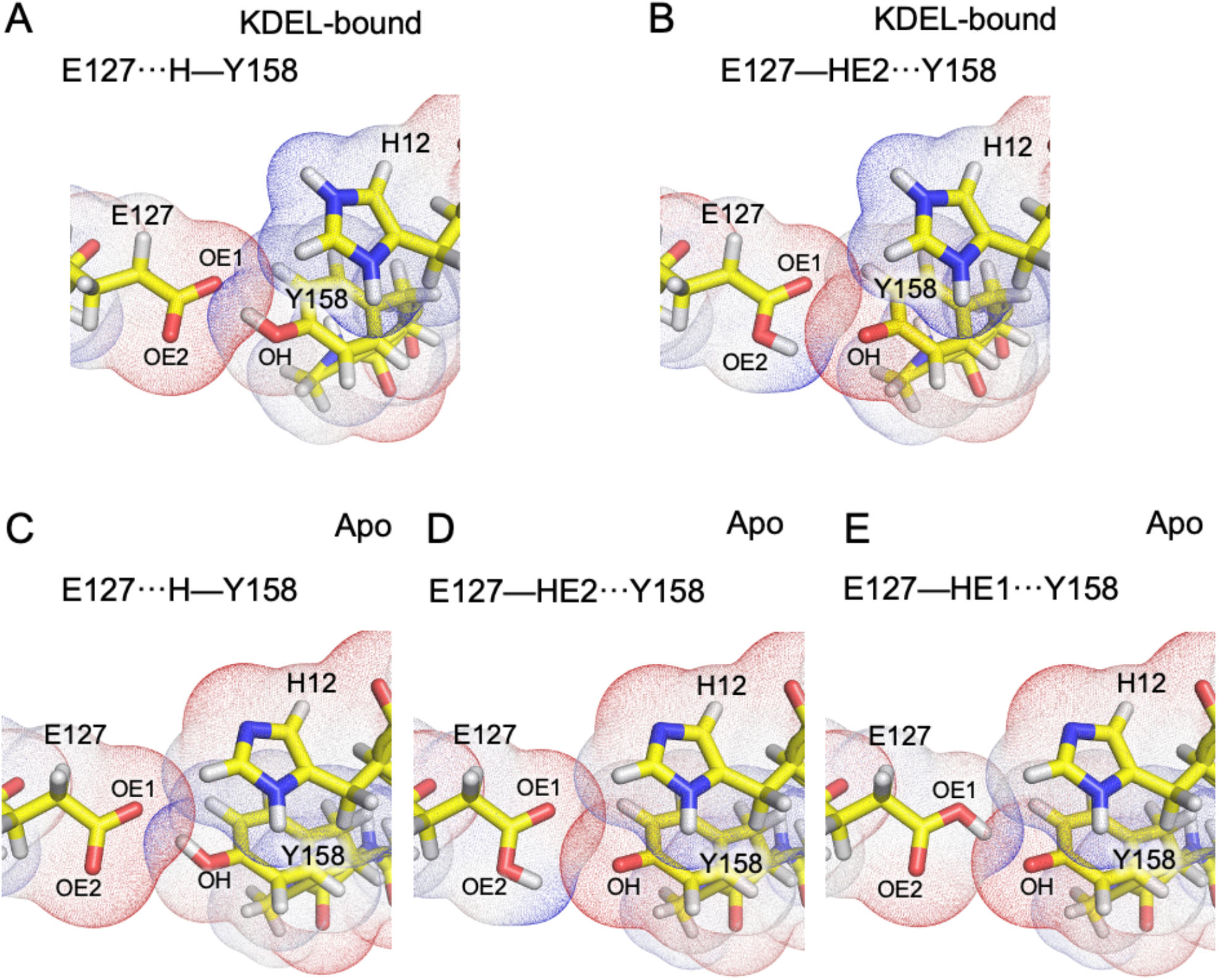
Electrostatic surface potentials. The region of negative electrostatic potential is given in red, and the region of positive electrostatic potential is given in blue. In the KDEL-bound conformation (A, B), E127···H—Y158 has complementary charge overlap during the region of the hydrogen bond (A), whereas the region of overlap in the E127—HE2···Y158 system is comprised mainly of like (negative) charge (B). Favourable overlap is also observed in E127···H—Y158 in Apo conformation for both the E127···H—Y158 system (C) and the E127—HE1···Y158 system but not the E127—HE2···Y158 (D) system.

Thus, all calculations are consistent with the expectation that the most favourable position for the proton in all states is close to the tyrosine oxygen.

### The strength of the E127···H—Y158 H-bond depends on the protonation state of H12

The proton position calculations suggested that in the KDEL-bound state, the energy barrier for the hydrogen to move to/from the nearest sidechain oxygen of E127 was likely to be very small. Given the potential role of H12 in mediating pH effects, we postulated that the protonation state of H12 might affect the strength of the E127···H—Y158 H-bond, which in turn would be expected to influence the ability of the receptor to lock the KDEL receptor in the activated state. We explored this aspect by examining the change in free energy required to transition from the KDEL-bound state to the Apo state under different protonation states of H12.

As is shown in **Table 2**, the free energy difference of transition from KDEL-bound to Apo state is highest when H12 is protonated (HIP: −15.83 - (−31.37)= 15.54 kcal/mol) than the neutral form of histidine (HID: −20.55-(−30.33)= 9.78 kcal/mol; HIE: −17.4-(−27.30)= 9.9 kcal/mol). Thus, 5.7 kcal/mol more free energy is required for the transition when H12 is protonated

**Table 2.**
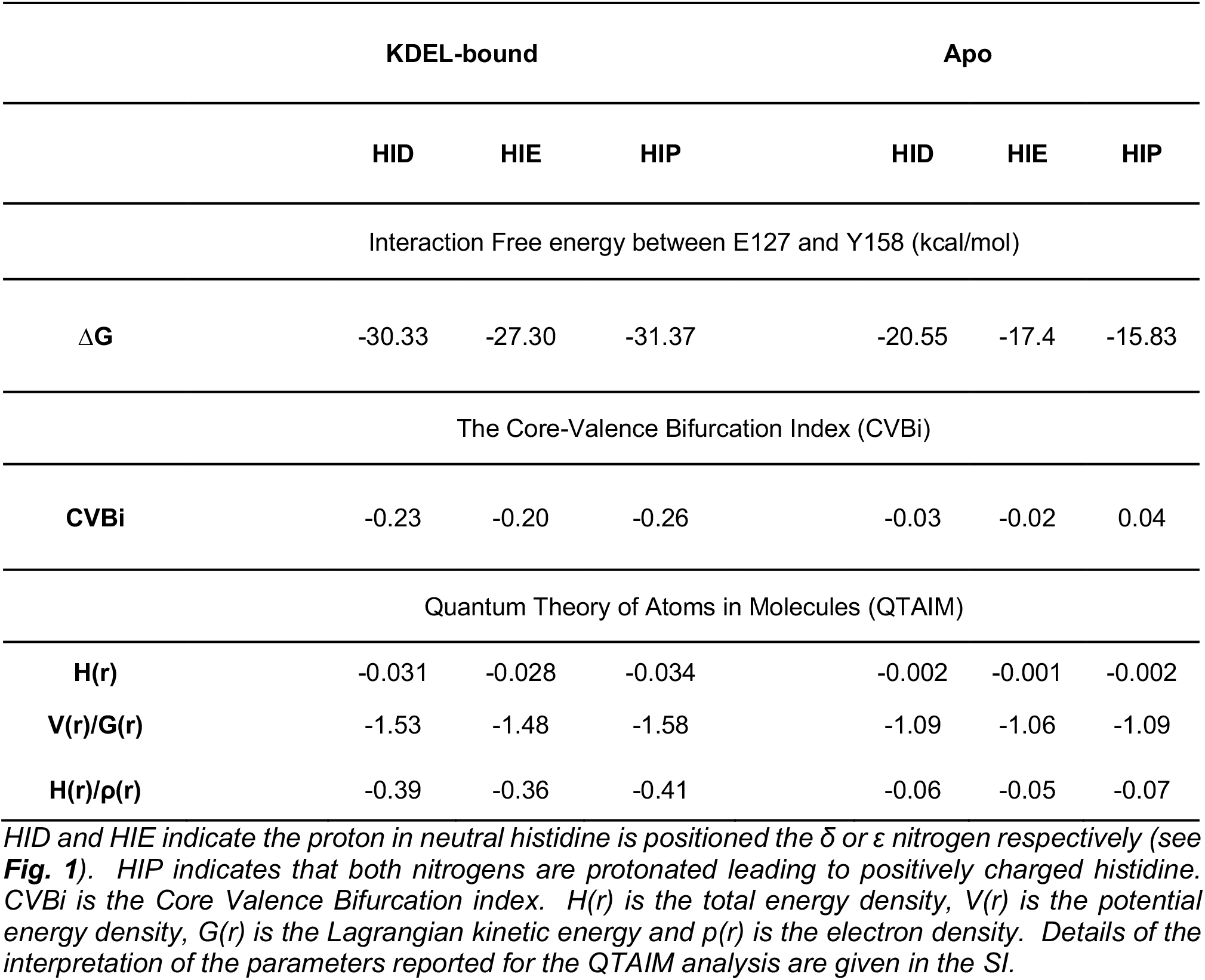
Properties of the E127···H—Y158 H-bond.

|  | KDEL-bound |  |  | Apo |  |  |
| --- | --- | --- | --- | --- | --- | --- |
|  | HID | HIE | HIP | HID | HIE | HIP |
| Interaction Free energy between E127 and Y158 (kcal/mol) |  |  |  |  |  |  |
| $\Delta G$ | -30.33 | -27.30 | -31.37 | -20.55 | -17.4 | -15.83 |
| The Core-Valence Bifurcation Index (CVBi) |  |  |  |  |  |  |
| CVBi | -0.23 | -0.20 | -0.26 | -0.03 | -0.02 | 0.04 |
| Quantum Theory of Atoms in Molecules (QTAIM) |  |  |  |  |  |  |
| H(r) | -0.031 | -0.028 | -0.034 | -0.002 | -0.001 | -0.002 |
| V(r)/G(r) | -1.53 | -1.48 | -1.58 | -1.09 | -1.06 | -1.09 |
| H(r)/$\rho(r)$ | -0.39 | -0.36 | -0.41 | -0.06 | -0.05 | -0.07 |

Having shown that the free energy penalty of transition is greatly enhanced when the histidine is protonated, we then assessed if the short hydrogen bond is influenced by the protonation state of H12. The crudest way of estimating H-bond strength is via the length of the bond between the hydrogen and hydrogen bond donor R(D-H) ^15^, as when the proton is shared more between the hydrogen bond donor and acceptor, it will stretch the R(D-H). As shown in **Table 2**, the R(D-H) distances in the KDEL-bound state are all longer than that in the Apo state, suggesting a stronger hydrogen bond in the KDEL-bound state compared with the Apo state. Furthermore, in the KDEL-bound state, the R(D-H) is longest when H12 is protonated, suggesting that protonation strengthens the hydrogen bond. Such an effect is not observed in the Apo state, where the R(D-H) is the same regardless of the protonation state of the histidine.

A more accurate estimate of H-bond strength can be obtained via the QM methods known as Core-Valence Bifurcation Index (CVBi) ^16^ and quantum theory of atoms in molecules QTAIM ^17^ (**see Methods and SI**). All these methods give a consistent result that the H-bond is stronger in the KDEL-bound state compared with the Apo state. For example, the CVBi values are much more negative for the KDEL-bound state than the Apo, and the total energy density (H(r)), is more negative in the KDEL-bound than in the Apo state. More details on the interpretation of the QTAIM values is given in the SI. Furthermore, note that in the KDEL-bound state, this hydrogen bond is further enhanced by the protonation of the histidine. This effect is not seen in the Apo state, where the strength of the H-bond seems to be independent of the protonation state of H12.

Though we have demonstrated that the protonated H12 strengthens the interaction between E127 and Y158 via enhancement of the hydrogen bond, it is still unclear as to how the protonation state of histidine would affect the strength of the hydrogen bond. A very short H-bond, such as that observed in the KDEL-bound state, is indicative of a low-barrier H-bond. To test if this hydrogen bond is indeed a low-barrier hydrogen bond, we computed the potential energy landscape of transferring the proton from the hydrogen bond donor (Y158) to the hydrogen bond acceptor (E127). If we consider the Apo state first (**Fig. 3A**), this shows a typical profile whereby there is a large (~20 kcal/mol) energy barrier to transfer the proton from the Y158 to E127. Moreover, there is very little effect from the protonation of H12. The profile for the KDEL-bound state is very different (**Fig. 3B**) and the protonation state of H12 can have significant effect. When the histidine is deprotonated, the energy difference between the global energy minimum (proton bound to the donor) and the local energy minimum (proton bound to the acceptor) is very high (~8 kcal/mol for HIE and ~7 kcal/mol for HID). However, when the histidine is protonated (HIP), this energy difference shrinks to only 4 kcal/mol. Thus, in the KDEL-bound state, only when H12 is protonated, can the hydrogen bond be classified as a low-barrier hydrogen bond and is consistent with our hypothesis that the protonated histidine strengthens the hydrogen bond in the KDEL-bound state.

**Figure. 3.**
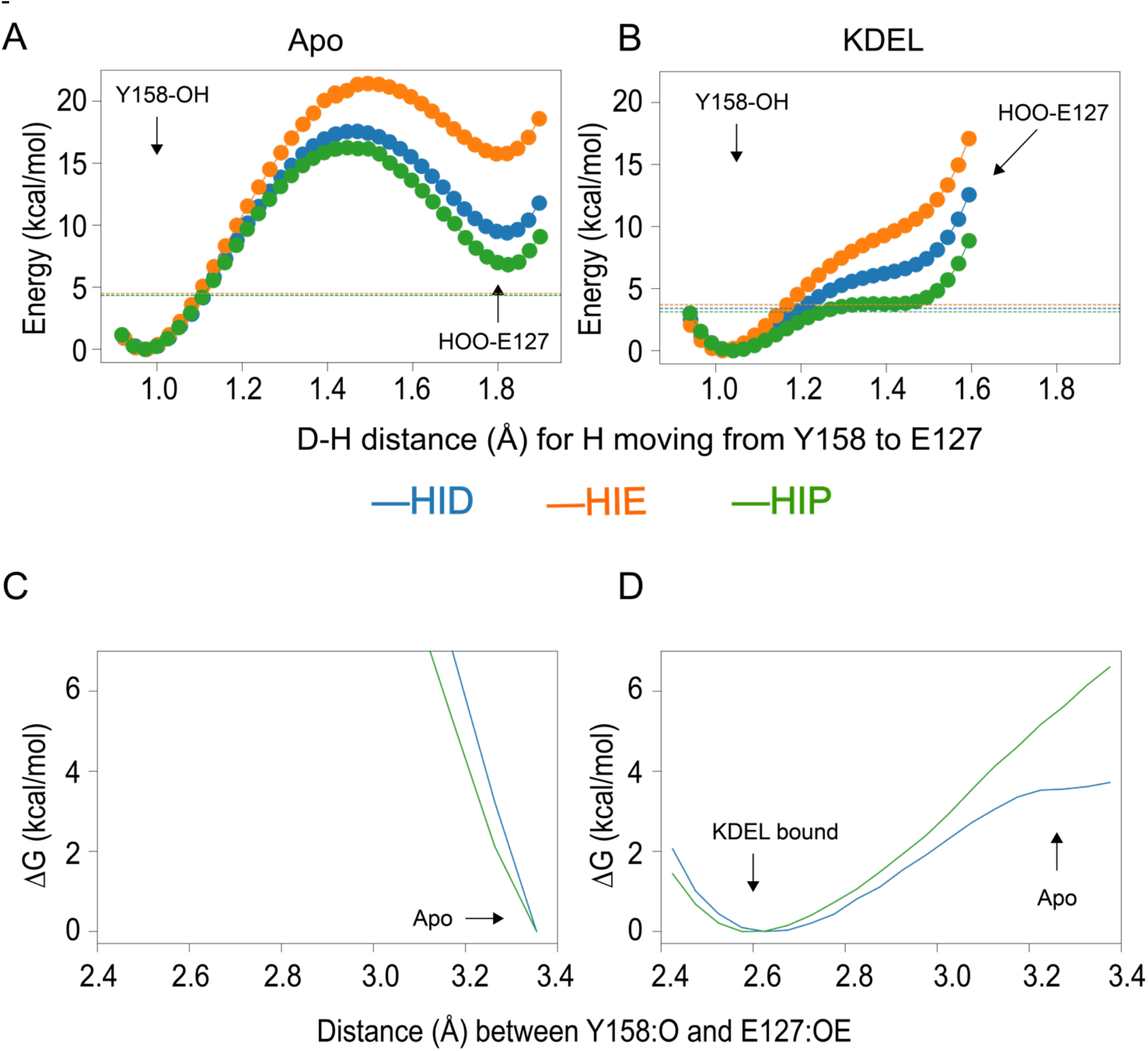
Free energy of proton transfer from the Y158 to E127. A. In the Apo state, a very large barrier (~20 kcal/mol) is observed during the proton transfer, which is not observed in the KDEL-bound state B. In the KDEL-bound state, the energy difference for the proton movement is smaller than the ZPE (dashed line) when the histidine is protonated (green). The energy difference is greatly increased when histidine is deprotonated (orange and blue). C. The energy landscape of separating the hydrogen bond. In the Apo state, the short hydrogen bond is prohibited by the larger energy barrier. D. In the KDEL-bound state, 6.5 kcal/mol of energy is required to break the hydrogen bond when histidine is protonated (green line), while only 3.5 kcal/mol is required when histidine is deprotonated (blue line).

Though the QM calculations showed that the protonated histidine allows the criteria of a low-barrier hydrogen bond to be met, a potential issue is that they are performed under vacuum conditions. Thus, to address this issue we also performed QM/MM calculations to compute the free energy barrier of stretching the hydrogen bond in both KDEL-bound and Apo states with both deprotonated histidine (HID) and protonated histidine (HIP). As shown in **Fig. 3C**, in the Apo state, the calculations demonstrate that the short hydrogen bond is unable to form regardless of the protonation state of the histidine. In the KDEL-bound state (**Fig. 3D**) however, when the histidine is deprotonated (HID), some initial resistance is met when separating the H-bond, which quickly plateaus resulting in only ~3.5 kcal/mol for the transition. However, when the histidine is protonated, the resistance is ever increasing, and the free energy penalty is around 6.5 kcal/mol when reaching the same distance as that observed in the Apo state.

### The Short Hydrogen Bond Influences the Affinity of H12 for Protons

Since protonated histidine stabilised the low-barrier hydrogen bond in the KDEL-bound state but had no effect on the hydrogen bond in the apo state, the reciprocal effect (i.e that the short hydrogen bond in the KDEL-bound state should stabilize a protonated H12, but no such effect will be present in the Apo state), should also be true. We employed a double bilayer approach (**Methods** and **Fig. 4**) to perform alchemical transformations simultaneously in the Apo and KDEL-bound states to compute the difference in proton affinity in these different states under two different regimens; one where the short hydrogen bond is imposed via harmonic restraint throughout the whole process and the other where there are no restraints to maintain the short H-bond distance (see **Methods**). The calculations show that under the conditions of the short H-bond, the protonation is more favourable in the KDEL-bound state compared to the apo state, by ~ 3.5 kcal/mol (**Table 3**). However, when the short H-bond is not constrained, this difference is reduced to ~2 kcal/mol (**Table 3**). Control calculations performed on histidines elsewhere on the protein (H11 and H90) suggest that any local variation of protonation is likely to be small, of the order of 0.5 kcal/mol.

**Table 3.** ΔΔG of (in kcal/mol) of deprotonation* of H12 in the KDEL-bound and protonation** of H12 in apo state under different conditions.

| Condition | Run 1 | Run 2 | Run 3 |
| --- | --- | --- | --- |
| Short H-bond imposed | 3.555 | 3.787 | 3.156 |
| No H-bond restraint | 1.992 | 1.996 | 2.176 |
| H90 (control – no restraint) | 0.847 |  |  |
| H11 (control – no restraint) | 0.435 |  |  |

\*to or from\*\* a neutral histidine with the proton on the $\delta$ carbon (HID).

**Figure 4.**
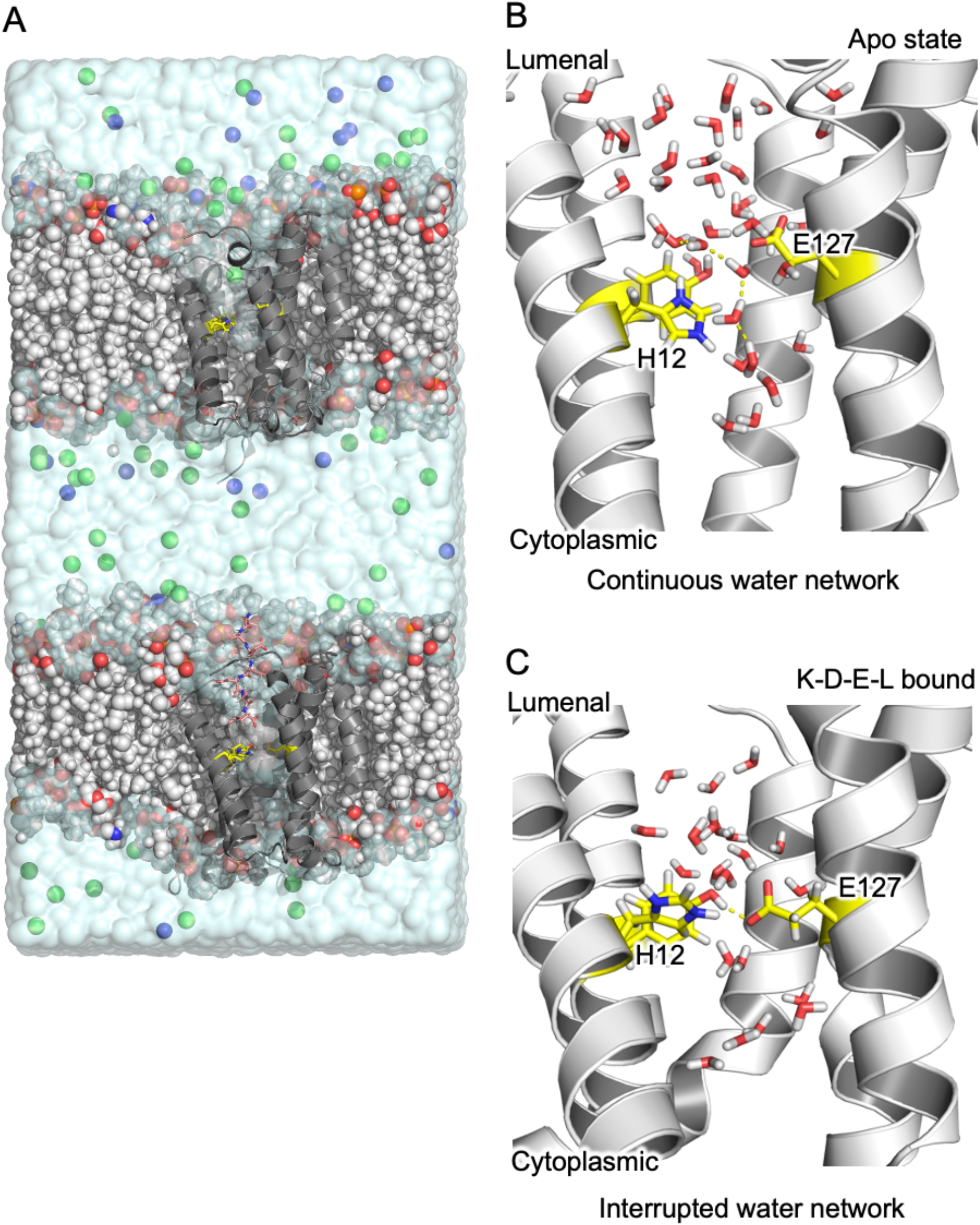
Influence of the short H-bond on H12 affinity. (A) shows the simulation box used, which is a double bilayer system to simultaneously add/remove a proton from H12 in each KDEL receptor. Na^+^ and Cl^−^ ions are represented as blue and green spheres respectively and the lipid bilayer is drawn as vdW spheres. (B) In the Apo state and when the H12 is protonated, a continuous water network linking the lumenal and cytoplasmic pockets is observed as highlighted by yellow dashed lines. (C) In the KDEL-bound state, the presence of the short hydrogen bond prohibits the formation of the water network. H12, E127 and Y158 are drawn as yellow sticks in all figure parts.

To further understand the molecular mechanism behind this short hydrogen bond mediated protonation event, we examined the local water movement around the short hydrogen bond. Topologically, the KDEL receptor resembles a transporter with both extracellular and intracellular pocket ^13^. The short hydrogen bond effectively acts as a gate between the extracellular and intracellular pocket in the receptor. In the Apo state, the two water pockets are joined by a water path due to the absence of the short hydrogen bond (**Fig 4B**). In the KDEL-bound state, on the other hand, the communication between two water pockets is prohibited by the short hydrogen bond (**Fig 4C**). This difference in local water dynamics gives rise to the ~3.5 kcal/mol free energy difference. In the absence of the short hydrogen bond, a sporadic water wire was observed to form in the KDEL-bound state resembling the local water environment in the Apo state, which brings the free energy difference down to ~2 kcal/mol. All of these calculations are consistent with our hypothesis that the protonated histidine strengthens the hydrogen bond and with the QM/MM calculations, where the transition from KDEL-bound state to apo state is 3 kcal/mol less when H12 is protonated.

## Discussion

In this work, we have shown that the distance between E127 and Y158 in the KDEL-bound crystal structure is consistent with what might be expected for a short H-bond. Short H-bonds, where the donor−acceptor distance is typically shorter than 2.7 Å are often found in the active sites of enzymes ^18^. In that context, such bonds have been described as “low-barrier H-bonds” ^19,20^. A feature of such bonds is that the proton is more readily delocalized between the donor and acceptor atoms and the resulting H-bond is predicted to be about 10-20 kcal/mol stronger than normal H-bonds ^21,22^. The results here confirm the initial postulation that in the KDEL-bound state there is an unusually short H-bond between E127 and Y158. The fact that it is a tyrosine is also rather interesting as a recent study revealed an unexpected enhancement of tyrosine in such interactions ^23^. The (QM) potential energy surface of proton transfer (**Fig. 3**) confirms that in the apo state there is a large and expected barrier to move the proton away from its preferred position. In the KDEL-bound state, the profile adopts a single well with a shoulder. At this distance, (2.5 Å between donor and acceptor oxygens), the result is entirely consistent with other reports of short H-bond proton potential energy surfaces (review recently by ^24^). Furthermore, the calculations suggest the ease of proton-sharing is significantly increased if H12 is protonated. H12, which has already been identified as the proton sensor ^13^, would be expected to be protonated in the Golgi, where it also binds to KDEL-bearing cargo proteins.

The above calculations suggest that the presence (and strength) of the short H-bond will be influenced by the protonation state of H12. We also investigated whether presence of absence of a short H-bond will influence the likely protonation state of H12. The results confirm that the presence of the short H-bond favours protonation of H12. Moreover, this appears to be linked to the dynamics of the surrounding water molecules (**Fig. 4B,C**). Inspection of trajectories suggests in the KDEL-bound state with a short H-bond present and H12 protonated, the water pockets either side of the H-bond are not connected. However, in all other states the water pockets are connected by a continuous h-bond network.

What does all this mean for the function of the receptor? The receptor binds KDEL-tagged proteins in the Golgi where the pH is lower than in the ER (where the receptor releases its cargo). H12 has already been proposed to be the pH sensor ^13^ and our observation here that a protonated H12 leads to a strengthening of the short H-bond suggests that it does play a role in forming a tighter complex with the ERS. At neutral pH (as found in the ER), the short H-bond is weakened, and this facilitates the release of the KDEL-containing cargo protein. We also know from previous structural work, that in the KDEL-bound state, there is a different conformation of helix 7 which moves I147 out of the cytoplasmic pocket. Based on our observations of the hydrogen-bonded networks that connect the two water pockets either side of H12, we postulate that the protonation state of the receptor may also play a role in controlling the water-dynamics, which in turn might control the preferred conformational state of the receptor. Clearly much more work is required to properly address this hypothesis. However, a mechanistic link between proton binding, water network rearrangements and structural stability are similar themes in proton coupled secondary active transporters ^25–27^ with which the KDELR shares the same topology and suggested evolutionary ancestry ^28^. We summarize our results in **Fig. 5**.

**Figure 5.**
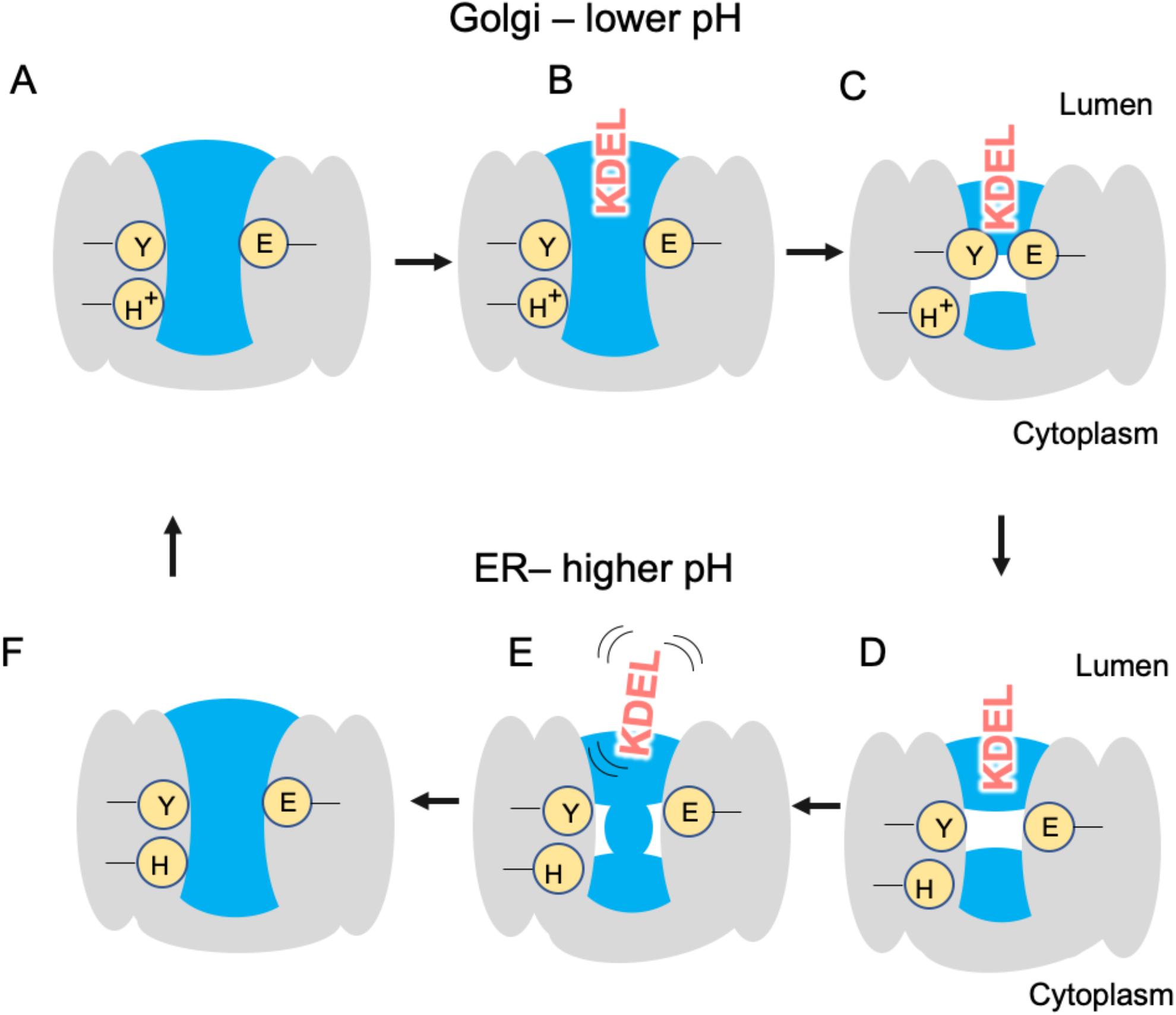
In the Golgi, the KDEL receptor is initially in its apo state, with the key histidine protonated and water present on both sides of E127/Y158. The KDEL-tagged substrate binds the receptor starting to trap waters as it enters (B). As binding completes some waters are cut from access to the lumen, the short hydrogen bond forms and the receptor undergoes other conformational changes to adopt the crystallographically observed KDEL-bound conformation (C). The KDEL receptor/KDEL-tagged substrate complex is then transferred to the ER where the higher pH causes deprotonation of the histidine. Without the protonated histidine, the short hydrogen bond is no longer maintained (D) and, the KDEL-tagged substrate is destabilized and dissociates (E). Since deprotonated histidine also favours a continuous water network, the receptor changes conformation and the water pocket is connected to bulk again (F).

## Conclusion

In this work we have performed extensive calculations that support the existence of a short H-bond in the KDEL receptor. Moreover, we have shown how the protonation state of a key histidine (H12) can dramatically affect the strength of that hydrogen bond. Via investigation of the reciprocal effect (of the short H-bond on the affinity of H12 for a proton) we also found a hydrogen-bonded water network that is broken in the presence of the short H-bond in the KDEL-bound state only and may have ramifications for the conformational dynamics of the receptor, something that we are currently exploring.

## Acknowledgements

We thank Irfan Alibay and Joanne Parker for useful discussions. We thank the Advanced Research Computing facility, the ARCHER UK National Supercomputing Services for computer time granted via the High-End Computing Consortium for Biomolecular Simulation, HECBioSim (http://www.hecbiosim.ac.uk), supported by EPSRC (EP/L000253/1). This research was supported by Wellcome Trust awards (109133/Z/15/A and 219531/Z/19/Z) to PCB and SN.

## Author contributions

SN and PCB designed the research. ZW designed, performed and analysed simulations. SN, ZW and PCB wrote the paper.

## Data availability

The authors declare no competing interests.

## Methods

### QM/MM simulations

The KDEL-bound KDEL receptor (KDELR) and the Apo KDEL receptor were retrieved from the PDB (Apo: 6I6B; KDEL-bound: 6I6H) and embedded in a DMPC bilayer following a previously described protocol ^29^. The QM region was defined as the three key residues (H12, Y158 and E127) and was described at the level of B97-3c, ^30^ whereas the qmcut was set to 3 nm to cover the whole protein. The force field amber 14SB ^31^ was used to describe the protein and LIPID17 ^32^ was used to describe the lipid. The simulation was run with ambertools19 ^33^ in conjunction with ORCA 4.2.0 ^34^. For the free energy calculation, the collective variable chosen was the distance between the oxygen in the hydroxyl group of the Y158 and the closet oxygen in the carboxyl group of E127. 11 equally spaced windows from 2.4 Å to 3.4 Å with a restraint of 25 kcal/mol/Å were run with a 1 fs timestep for 20 ps. The free energy profile was constructed with WHAM 2.0.9.1. ^35^.

### QM Geometry Optimisation

The coordinates of residues H12, E127 and Y158 were extracted from the crystal structure of both Apo (PDB: 6I6B) and the KDEL bound (PDB: 6I6H) KDELR. Note that in the original crystal structures the oxygen closest to the tyrosine OH is labelled OE2. To simplify the analysis here, we have relabelled that OE1 in this work. The N-termini of the three residues were capped with an acetyl group and the C-termini were capped with an N-methyl group. Hydrogen atoms were added to the crystal structure and different protonation states of the three residues were assigned using Maestro. The structure of the H12, E127 and Y158 complex, as well as each capped amino acid monomer and all the possible dimers (H12/E127; H12/Y158) were independently optimised with non-hydrogen atoms constrained using B3LYP together with the def2-TZVP ^36^ basis set with DFT-D4 ^37^ for the empirical dispersion correction. Due to a known bug in the ORCA frequency calculations, the following flags (%freq Hess2elFlags[0] 0 Hess2elFlags[3] 0 end) were added as is suggested by the developer (https://orcaforum.kofo.mpg.de/viewtopic.php?f=26&t=5566). Vibrational frequency analysis was done at the end of optimisation using ORCA 4.2.0 ^34^. The CVBi ^16^ and AIM analysis was performed with Multiwfn 3.7 ^38^.

### QM Interaction Energies

Single point energies were computed on the optimised structure with DLPNO-CCSD(T)/aug-cc-pVTZ ^39^ at the level of normal PNO, with aug-cc-pVTZ/JK for RIJK acceleration and aug-cc-pVTZ/C ^40^ for electron correlation calculations. The thermal correction for the Gibbs free energy was computed from the frequency calculations and corrected with quasi-harmonic correction ^41^ and a zero-point energy (ZPE) scaling factor of 0.985 ^42^ using Shermo ^43 to 310 K. For complexes with more than one amino acid, the energy is corrected for by basis set superposition error (BSSE). BSSE is defined as half {Burns, 2014 #9127^ of the BSSE energy. The interaction energy between E127(E) and Y158(Y) under the influence of the H12(H) is defined as

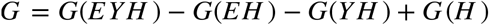

Where *G(EYH), G(EH), G(YH), G(H)* are the free energy of the geometry optimised E127, Y158, H12 complex, geometry optimised energy of the H12 in complex with E127 or Y158 and the energy of geometry optimised histidine, all with counterpoise corrected.

### QM Proton transfer energy

The energy of transferring a proton from tyrosine to glutamate was computed at the level of PWPB95-D4/def2-QZVPP//B3LYP-D4/def2-TZVP ^44^. The calculation was performed in the presence of three different protonation states of H12. During the geometry optimisation, the positions of all the non-hydrogen atoms were constrained and the bond between the proton and the closest oxygen was also constrained. The vibration mode corresponding to motion of the key proton was taken as the mode with the highest infrared intensity and confirmed with visual inspection using Gaussview 6.0. The ZPE corresponding to this mode was computed using Shermo ^43^ .

### Protonation free energy calculations

A double bilayer box with KDEL-bound and Apo protein was constructed. Bilayers were placed 3 nm apart and solvated with TIP3P water and ions added up to 150 mM NaCl. The short hydrogen bond was maintained with a harmonic restraint joining the two oxygens in E127 and Y158 with restraint strength of 10,000 kj/mol/nm. The equilibrium length is taken from the crystal structure. The topology for alchemical transformation was generated with pmx ^45^, where the histidine in the KDEL-bound structure was transformed from HIP to HID, while the same histidine in the Apo state were transformed from HID to HIP. 21 equally spaced windows were used during the transformation, where the bonded, charge and vdw parameters were changed linearly at the same time. The simulations were run with Gromacs 2019.1 ^46^, with parameters taken from ^47^. After the initial energy minimisation, a 200 ps simulation in the NVT ensemble were conducted, followed by a 1 ns run in the NPT ensemble. The production run was performed with replica exchange at an interval of 1 ps for 30 ns. The resulting energy files were analysed with alchemical exchange ^48^ with the first 5 ns discarded as equilibration time.

