## Supplementary Information for "The KDEL Trafficking Receptor Exploits pH to Tune the Strength of an Unusual Short Hydrogen Bond"

**Details on the Quantum Theory of Atoms in Molecules (QTAIM)**

According to Bader's the quantum theory of atoms in molecules (QTAIM) theory, topology analysis of the electron density, ($\rho_r$) and its Laplacian can be used to give a quantitative description of the hydrogen bond system. For a bond to present, an atomic bond path has to link the the two atoms that form the bond and a bond critical point (BCP) has to be present on this path[1,2]. In all hydrogen bond systems, an atomic bond path was found between the proton and the hydrogen bond acceptor and a BCP is present on this path. Table 2 of the main manuscript lists some important topological parameters for this BCP, including the total energy density H(r)[3], which is the sum of Lagrangian kinetic energy G(r) and potential energy density V(r)[4], the V(r)/G(r), H(r)/ρ(r), where ρ(r) is electron density. The strength of the hydrogen bond can be inferred from H(r), where Rozas et al[5] proposed that a weak hydrogen bond shows H(r) >= 0 and a strong hydrogen bond shows H(r) < 0.

The hydrogen bonds formed in the KDEL-bound states between the Y158H and E127 can be classified as a strong hydrogen bond while the hydrogen bonds in the apo state are classified as a weak hydrogen bond. The strongest hydrogen bond is observed when the histidine is protonated (HIP) in the KDEL-bound state. Espinosa et al.[4] purposed both H(r)/ρ(r) and V(r)/G(r) can serve as an indication of the bond strength. H(r)/ρ(r) is also commonly called the bond degree (BD) and a smaller value indicates a stronger bond. Similar to BD, the smaller the V(r)/G(r), the stronger the bond. The rank of hydrogen bond strength is consistent with all other analysis where the hydrogen bond is stronger in KDEL-bound state compared with the apo state and the hydrogen bond in the KDEL-bound state under protonated histidine (HIP) is the strongest.
